# Acetate and hypertonic stress stimulate organelle membrane fission using distinct phosphatidylinositol signals

**DOI:** 10.1101/398685

**Authors:** Dipti Patel, Christopher Leonard Brett

## Abstract

Organelle morphology reflects an equilibrium between membrane fusion and fission that determines size, shape and copy number. By studying the yeast vacuole as a model, the conserved molecular mechanisms responsible for organelle fusion have been revealed. However, a detailed understanding of vacuole fission and how these opposing processes respond to the cell cycle, osmoregulation or metabolism to change morphology remain elusive. Thus, herein we describe a new fluorometric assay to measure vacuole fission in vitro. For proof-of-concept, we use this assay to confirm that acetate, a key intermediary metabolite, triggers vacuole fission in vitro and show that it also blocks homotypic vacuole fusion. The basis of this effect is distinct from hypertonic stress, a known trigger of fission and inhibitor of fusion that inactivates the Rab-GTPase Ypt7: Treatment with the phosphatidylinositol-kinase inhibitor wortmannin or the catalytic domain of the Rab-GAP (GTPase Activating Protein) Gyp1 reveal that fission can be triggered by Ypt7 inactivation alone in absence of hypertonic stress, placing it upstream of PI-3,5-P_2_ synthesis and osmosis required for membrane scission. Whereas acetate seems to block PI-4-kinase to possibly increase the pool of PI on vacuole membranes needed to synthesize sufficient PI-3,5-P_2_ for fission. Thus, we speculate that both PI-4-P and PI-3-P arms of PI-P signaling drive changes in membrane fission and fusion responsible altering vacuole morphology in response to cellular metabolism or osmoregulation.

**GRAPHICAL ABSTRACT:** 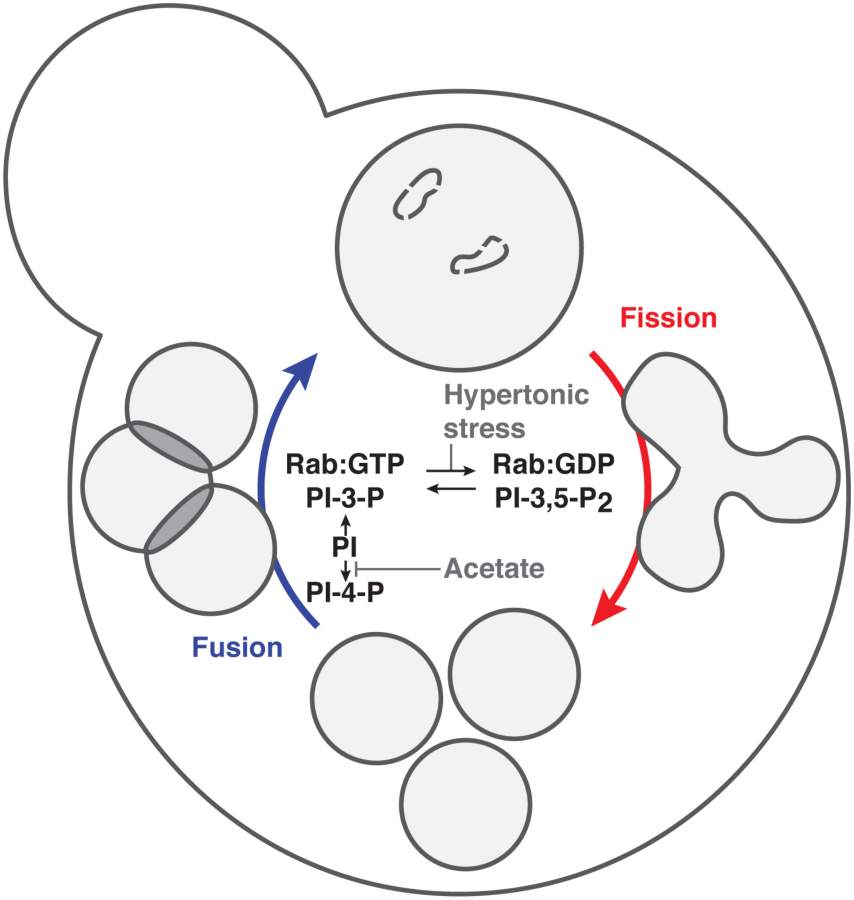

## INTRODUCTION

Morphology of most organelles is determined by membrane fusion and fission (also called fragmentation). These include mitochondria, chloroplasts, the Golgi apparatus, peroxisomes and organelles of the endocytic pathway including endosomes and lysosomes (or vacuoles in yeast; *Shorter and Warren, 2002; Weisman, 2003; Friedman and Nunari, 2014*; *Luzio et al., 2014; Knoblach and Rachubinski, 2016*). These opposing processes drive changes in organelle size, number and shape for cellular responses to environmental changes or signaling events or for organelle inheritance during cell division. Endosomes and lysosomes rely on cycles of fusion and fission for membrane trafficking for endocytosis (*Gautreau et al., 2014; Luzio et al., 2014*). Large numbers of endosomes or lysosomes generated through fission produce enough to be deposited throughout the cell as required for diverse functions, including cell signaling, plasma membrane repair and intra-organelle communication (*Pu et al., 2016; Cabukusta and Neefjes, 2018*). Enlargement of lysosomes and vacuoles rely on fusion to accommodate autophagy, synchronized with changes in amino acid metabolism by TOR (Target Of Rapamycin) signaling – a key mediator of cell metabolism particularly when cells are starved or growing (*Perera and Zoncu, 2016*).

Most knowledge of the molecular machinery underlying these processes has been gleaned by studying the budding yeast vacuole as a model. *Saccharomyces cerevisiae* cells typically contain 2 – 5 vacuoles that undergo regulated cycles of membrane fission and fusion (*Weisman, 2003; Li and Kane, 2009*). Because they are relatively large (0.5 – 3 µm diameter) and can be exclusively stained with many vital dyes (e.g. FM4-64), vacuole morphology is easily assessed by fluorescence microscopy (*Conibear and Stevens, 2002*). Vacuoles are easily purified permitting further biochemical study of organelle membrane fusion and fission in vitro (*Conradt et al., 1992*). Yeast is of course a genetically tractable model system, permitting genetic analysis as well (*Struhl, 1983*). Using this system, it was discovered that these processes are highly coordinated, as one process must dominate to effectively change and retain morphology, e.g. to increase copy number, fission is stimulated whilst fusion is blocked (*LaGrassa and Ungermann, 2005; Durchfort et al., 2012*). These findings have led to the idea that the underlying machinery is highly integrated (e.g. *LaGrassa and Ungermann, 2005; Alpadi et al., 2013*). The basis of homotypic vacuole fusion has been resolved with incredible molecular precision (see *Wickner, 2010*). However, vacuole or organelle fission is less understood.

Within live yeast cells, vacuoles fragment (i.e. undergo membrane fission) during the cell cycle and in response to hyperosmotic stress, oxidative stress, or TOR signaling stimulated by ER stress (*Bonangelino et al., 2002; Weisman, 2003; LaGrassa and Ungerman, 2005; Stauffer and Powers, 2015*). Through in vivo and in vitro analysis, it was shown that vacuole fission is a two-step asymmetrical process that requires phosphoinositol-3,5-diphosphate (PI-3,5-P_2_) generated from phosphoinositol-3-phosphate (PI-3-P) on the cytoplasmic face of the vacuole lipid bilayer by Fab1, a PI-3-P5 kinase (or PIKfyve in mammals), in complex with Vac14, Vac7 and Fig4 (*Dove et al., 1997; Bonangelino et al., 2002; Michaillat et al., 2012; Zieger and Mayer, 2012*). Also implicated in this process is the H^+^-electrochemical gradient maintained by the V-type H^+^-ATPase (*Baars et al., 2007; Bonangelino et al., 2002; Michaillat et al., 2012; Stauffer and Powers, 2015*), which likely occurs downstream of Fab1 as its activity is supported by PI-3,5-P_2_ (*Li et al., 2014; Ho et al., 2015*). The process is thought to culminate with lipid bilayer scission by the dynamin-like GTPase Vps1 in coordination with the PROPPIN Atg18, which binds to the Fab1-complex and responds to PI-3,5-P_2_ (*Peters et al., 2004; Baars et al., 2007; Efe et al., 2007; Takeda et al., 2008; Gopaldass et al., 2017*).

A decrease in lumenal volume is also necessary to accommodate perimeter membrane collapse and constriction at sites of scission. Currently, it is not entirely how this occurs, but insight has been gleaned by experimentally applying hypertonic stress, to drive water out of the vacuole lumen. From these studies, it was revealed that Vac14 is required for activation of Fab1 in response to a decrease in organelle volume induced by hypertonic stress (*Bonangelino et al., 2002*). Recently, Ivy1, an inhibitor of Fab1, inverted BAR (I-BAR) protein and effector of the Rab-GTPase Ypt7 was also implicated in this response (*Malia et al., 2018*): When Ypt7 is inactivated by hypertonic stress (*Brett and Merz, 2008*), fusion is halted and the Rab disengages Ivy1. By also possibly sensing a change in membrane lipid packing or lateral tension induced by loss of lumenal volume, Ivy1 then releases Fab1, activating it to generate PI-3,5-P_2_ and drive fission. Thus, the coordination of PI signaling and Rab activity are key modulators of vacuole fission and fusion. However, there are many outstanding questions related to how these molecular mechanisms interact to drive changes in both organelle membrane surface area and lumenal volume required for fission.

For example, given the newfound role for Ivy and Ypt7 in stimulating fission, is inactivation of Ypt7 by a Rab-GAP (Rab-GTPase Activating Protein) sufficient to drive fission in absence of hypertonic stress? Or is it needed to induce changes in membrane properties to inactivate Ivy1 (see *Malia et al., 2018*)? Also, acetate was shown to stimulate vacuole fission in vitro (*Michaillat et al., 2012*). It was suggested that the underlying mechanisms that respond to acetate were unrelated to hypertonic stress but this was not formally tested. Furthermore, the question remains: how does acetate stimulate vacuole fission?

Herein we developed a new quantitative in vitro vacuole membrane fission assay and used PI-3-kinase inhibitor wortmannin and the recombinant Rab-GAP protein rGyp1-46 that targeting PI and Rab signaling, respectively, to answer these questions and refine our understanding of vacuole membrane fission and organelle morphology.

## RESULTS AND DISCUSSION

### A new assay to measure vacuole membrane fission in vitro

Until now researchers have relied on a fluorescence microscopy-based, semi-quantitative assay to estimate the number of vacuole fission products formed in vitro: The number of BODIPY FL-DHPE–stained small (< 0.6 µm diameter), medium (0.6 – 1.5 µm) and large (≥ 1.5 µm) vacuoles found in images of vacuole fission reactions were counted and the fraction of small vacuoles was calculated and reported as a fragmentation index (*Michaillat et al., 2012*). As an alternative, we designed a new simple, quantitative in vitro vacuole membrane fission assay (Figure 1A). It involves separating products of fission from larger vacuole precursors using low-speed differential centrifugation, whereby larger more dense vacuoles sediment and less dense smaller vacuoles remain suspended in the fission reaction buffer. To eliminate the need to stain vacuole membranes for detection (with FM4-64 or BODIPY FL-DHPE for example), we isolated vacuoles from yeast cells expressing Vph1, the stalk domain of the V-type H^+^-ATPase, tagged with GFP at its C-terminus. Vph1-GFP was used because it is known to be uniformly distributed on vacuole membranes (*Wang et al., 2002; McNally et al., 2017*) and thus should decorate fission precursor and product membranes at equal density. Using a plate-reading fluorometer, we then measured GFP fluorescence in the supernatant and pellet and report the ratio of background-subtracted supernatant fluorescence over total fluorescence (recorded from the supernatant and pellet) as a measure of vacuole fission in vitro.

**Figure 1.**
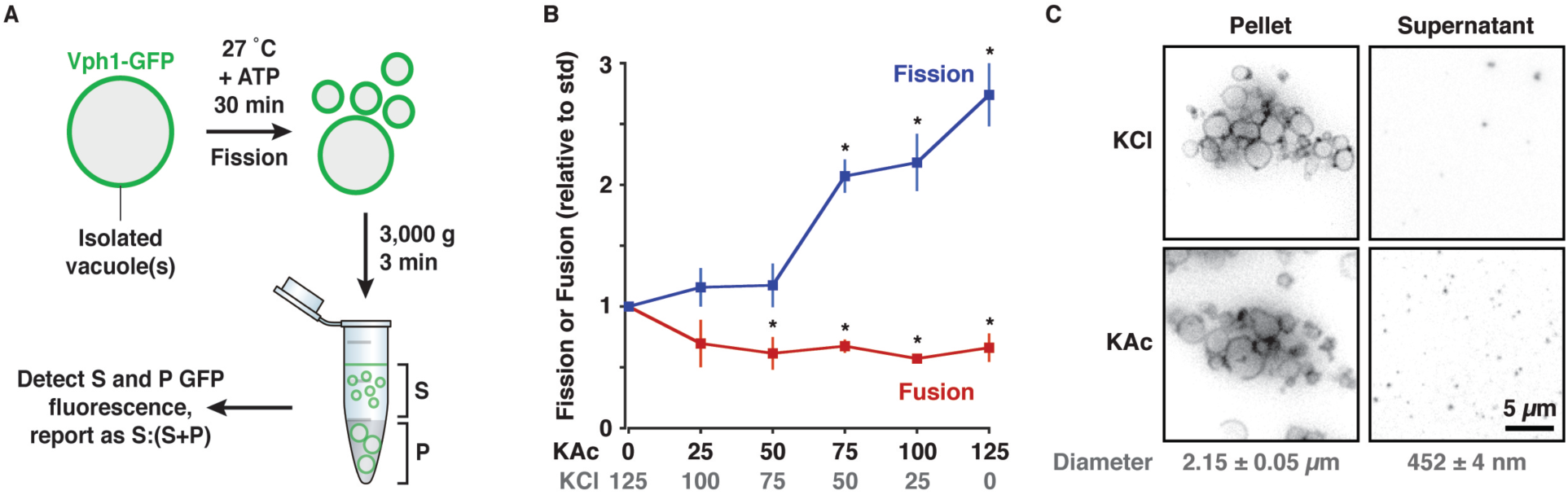
A new, simple cell-free vacuole membrane fission assay. (A) Cartoon depicting new fluorometric in vitro vacuole membrane fission assay. S, supernatant; P, Pellet. (B) Homotypic fusion or fission of isolated vacuoles in the presence of increasing concentrations of KAc in place of KCl. Means ± S.E.M. shown and P < 0.05 (*) as compared to standard conditions (125 mM KCl). (A) Micrographs of fission reactions containing vacuoles isolated from Vph1-GFP expressing yeast cells conducted in the presence of 125 mM KAc or KCl. Supernatants (containing fission products) and pellets (containing large vacuoles) are shown. Means ± S.E.M of vacuole diameters measured by quasi-elastic light scattering are indicated for each fraction.

To test this new method, we first conducted the assay under conditions previously shown to optimally drive vacuole fission in vitro, i.e. incubation at 27 °C for 30 minutes, ATP added as an energy source, and KAc added in place of KCl (*Michaillat et al., 2012*). As expected, we found that replacing KCl with KAc stimulates vacuole fission in vitro using this new assay (Figure 1B), whereby complete KAc replacement showed a significant 2.71-fold increase in fission, similar to that previously reported using the microscopy-based assay (*see Figure 2B in Machaillat et al., 2012*). However, in contrast to previous results, purified cytosol was not required for equally robust vacuole fission in our hands, suggesting that the underlying machinery co-purifies with the organelles. As fusion and fission machinery is highly integrated, we hypothesized that KAc should also inhibit fusion. As expected, using a lumenal content mixing assay, we found that homotypic vacuole fusion was inhibited by KAc replacement in vitro (Figure 1B), confirming that when one process is stimulated, the opposing process is blocked.

**Figure 2.**
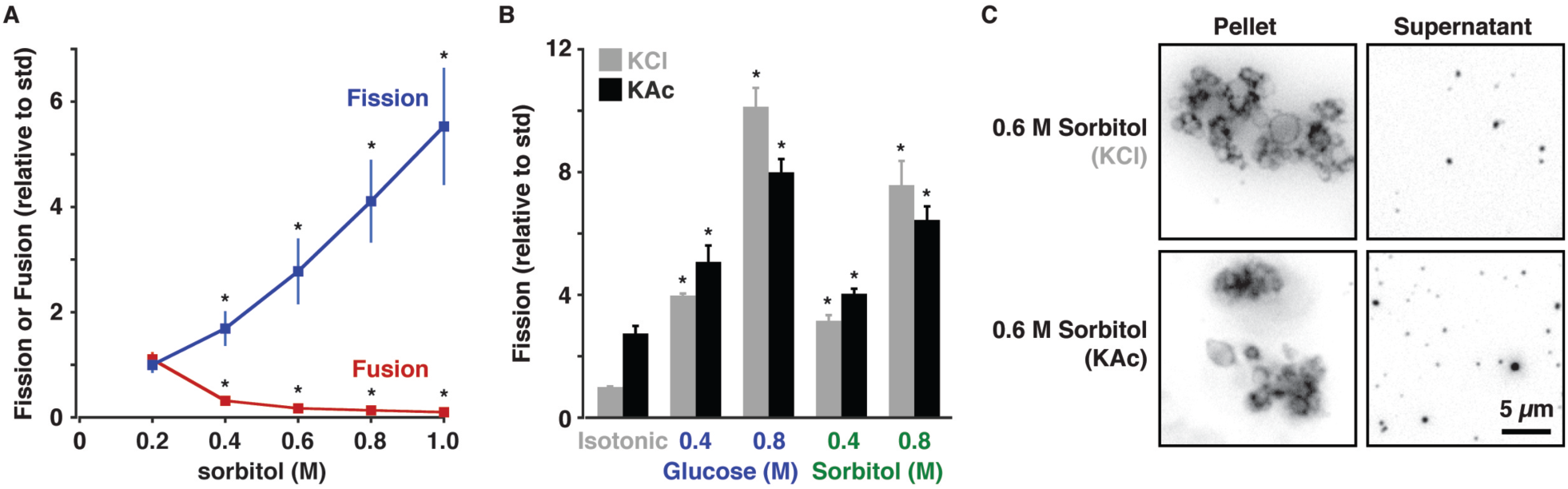
Effects of KAc and hypertonic stress on vacuole fission are additive. (A) Homotypic fusion or fission of isolated vacuoles in the presence of increasing concentrations of sorbitol. (B) Fission of isolated vacuoles in the presence of either glucose or sorbitol and 125 mM KCl or KAc. Means ± S.E.M. shown and P < 0.05 (*) as compared to standard, isotonic conditions (125 mM KCl, 200 mM sorbitol). (C) Micrographs of fission reactions containing vacuoles isolated from Vph1-GFP expressing yeast cells conducted under hypertonic conditions (0.6 M sorbitol) in the presence of 125 mM KAc or KCl. Supernatants (containing fission products) and pellets (containing large vacuoles) are shown.

To verify that we separated smaller fission products from larger precursors using this method, we imaged pellets and supernatants by HILO microscopy in the presence of 125 mM KCl, when fission is inhibited, or 125 mM KAc when fission is stimulated (Figure 1C). As expected, larger vacuoles were only observed in the pellet and smaller vesicles were observed in the supernatant. This was confirmed by measuring vacuole size by quasi-elastic light scattering, whereby vacuole diameter was 4.6 times smaller in the supernatant than in the pellet (Figure 1C). Of note, the diameter of the fission products (0.452 ± 004 µm) was consistent with a previous estimate (0.45 ± 0.27 µm) obtained from electron micrographs of vacuole fission reactions (*Machaillat et al., 2012*). Furthermore, when we counted puncta using micrographs of these samples, we found 2.84 ± 0.16 (n ≥ 7) times more GFP-positive vesicles in the supernatant of reactions conducted in the presence of KAc as compared to KCl, consistent with data acquired by fluorometry (Figure 1B). Thus, we are confident that this new assay is a valid method to accurately measure vacuole membrane fission in vitro.

### Effects of hyperosmotic shock and KAc on vacuole fission are additive

Exposure to hypertonic stress stimulates vacuole fission in live yeast cells and to remain fragmented, vacuole fusion is also inhibited (*LaGrassa and Ungermann, 2005*). The latter was confirmed in vitro, whereby treating vacuole fusion reactions with increasing concentrations of the osmolyte sorbitol blocked fusion in vitro (*Brett and Merz, 2008*). However, the effects hypertonic stress caused by sorbitol or other osmolytes on vacuole fission have not been extensively investigated in vitro. Thus, we examined the effect of adding sorbitol on vacuole fission using this new assay and found that, as expected, fission increases proportionally with sorbitol concentration (Figure 2A), whereby 1 M sorbitol shows a 5.6-fold increase in fission as compared to isotonic conditions (200 mM sorbitol, 125 mM KCl). As a control, we also confirmed that homotypic vacuole fusion is inhibited by increasing sorbitol concentrations (Figure 2A), confirming that these opposing processes are inversely regulated by hypertonic stress. It also reveals that the hypertonic stress directly affects fission machinery on the vacuole, and this response is not dependent on other mechanisms implicated in the yeast cell response to osmotic stress absent from the vacuole preparation, e.g. the Hog1 signaling machinery found in the cytoplasm and plasma membrane (*Brewster and Gustin, 2014*).

Next, to determine if KAc and sorbitol target the same underlying fission machinery, we examined the effect of adding both stimuli together. We hypothesized that if they target distinct machinery, then effects of each stimulus should be additive. Whereas if they target the same machinery, adding sorbitol to KAc should not induce a further increase in fission. We found that addition of sorbitol to buffer containing 125 mM KAc in place of KCl further stimulated the fission reaction (Figure 2B), suggesting that the stimuli were additive. To demonstrate that this effect was caused by hypertonic stress, as opposed to other chemical properties of sorbitol, we repeated the experiment with glucose, a different osmolyte, and obtained a similar result, although equimolar concentrations elicited significantly stronger responses in the presence of KCl or KAc (Figure 2B). To confirm that these conditions were indeed inducing fission and not causing lysis or somehow permitting larger vacuoles to contaminate the supernatant, we imaged the fission reactions by HILO microscopy (Figure 2C). These micrographs confirmed that only small vesicles were present in the supernatant fraction. Thus, we concluded that hypertonic stress and acetate trigger vacuole fission by independent mechanisms.

### KAc may inhibit a PI4-kinase to promote vacuole fission

Previously, acetate was shown to stimulates vacuole membrane fission in vitro but it remains unclear how it triggers this process (*Machaillat et al., 2012*). Although PI-3-P and PI-3,5-P_2_ are critical for both fusion and fission respectively, little attention has been given to the potential roles for PI-4-P and PI-4,5-P_2_ in fission. It is likely that they contribute because both have been implicated in vacuole fusion in vitro (*Stroupe et al., 2006; Mima and Wickner, 2009*). Furthermore, deleting genes encoding the enzymes responsible for their synthesis (STT4 or MSS4) cause vacuole morphology defects in vivo (*Audhya et al., 2000*). Thus, given that hypertonic shock targets PI-3-P and PI-3,5-P_2_ signaling and acetate likely targets another mechanism, we hypothesized that acetate may target PI-4-P and/or PI-4,5-P_2_ biosynthesis to trigger fission.

To test, this hypothesis we acutely inhibited PI-4-P synthesis in vitro using the PI-kinase inhibitor wortmannin. Although it blocks mammalian PI3-kinase activity, the yeast type III PI3-kinase Vps34 (the only PI3-kinase in *S. cerevisiae*) is insensitive to this drug (*Stack and Emr, 1994*). Rather, it has been reported to block PI-4-P synthesis by the type II PI4-kinase Stt4 (*Cutler et al., 1997*), and is thought to target orthologous PI4-kinases including Lsb6 found on vacuole membranes (*Han et al., 2002*). This inhibitor was used instead of a genetic approach, i.e. knocking out STT4, because of anticipated pleitropic effects given that PI-4-P and PI-4,5-P_2_ are important lipids for signaling at the plasma membrane and influence other vacuole functions, e.g. TOR signaling and autophagy (*Audhya et al., 2000; Tabuchi et al., 2006; Garrenton et al., 2010; Wang et al., 2012*). We find that increasing concentrations of wortmannin have no effect on fission stimulated by KAc (Figure 3A), suggesting that perhaps a PI4-kinase is already inhibited under these conditions rendering wortmannin ineffective. Importantly, we show that the concentrations of wortmannin used are bioactive as it further stimulated fission in the presence of glucose and KCl (Figure 3A). This important result suggests that (1) inhibition of PI-4-P synthesis can promote vacuole fission, and (2) hypertonic stress does not target PI-4-P production to induce fission, consistent with previous reports (e.g. *Bonagelino et al., 2002*). Thus, we conclude that acetate and hypertonic stress likely alter production of different PI-P species to stimulate vacuole fission.

**Figure 3.**
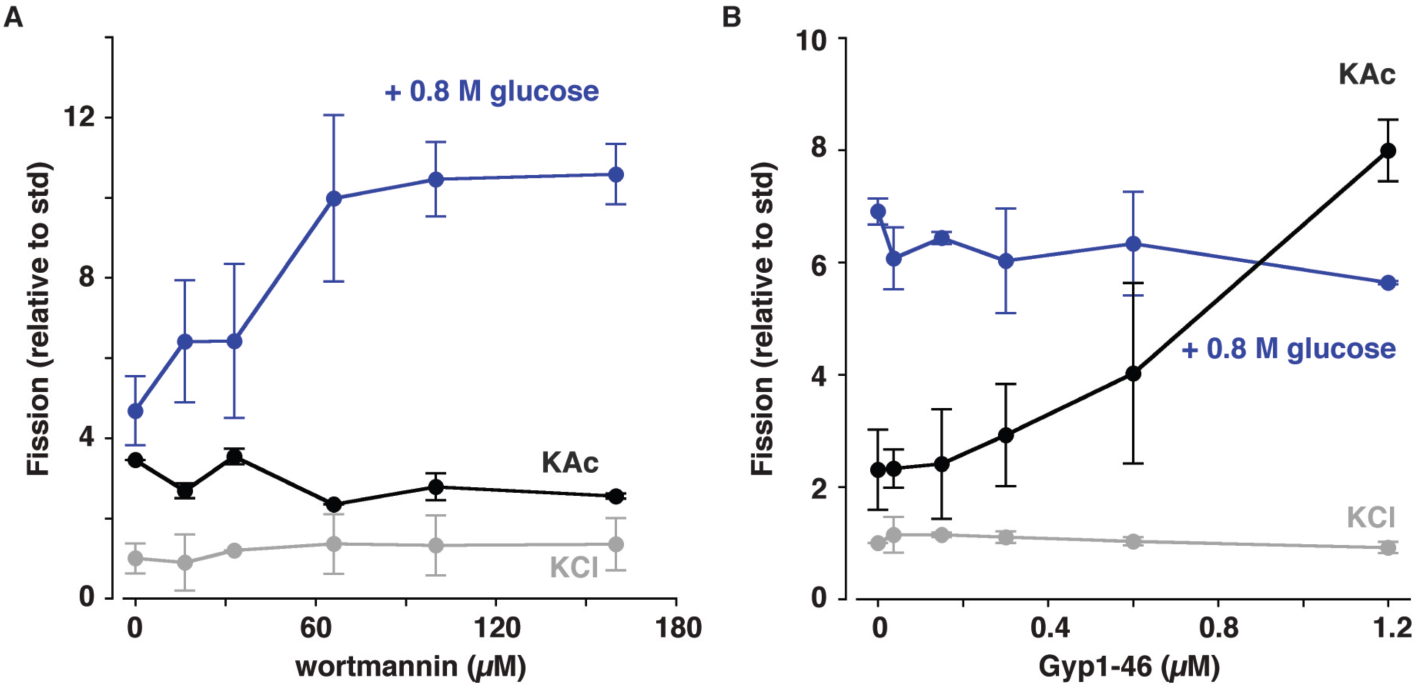
Effects of wortmannin or rGyp1-46 on vacuole fission triggered by acetate or hypertonic stress. Fission of isolated vacuoles in the presence of increasing concentrations of (A) the PI3-kinase inhibitor wortmannin or (B) the Rab-GTPase inhibitor rGyp1-46. Fission reactions contained either 125 mM KAc in place of KCl or 125 mM KCl with 0.8 M glucose. Means ± S.E.M. shown.

Why does acetate promote vacuole fission? Acetate, when ligated to coenzyme A, becomes acetyl-CoA, a central player in intermediary metabolism that facilitates macromolecular (e.g. fatty acid, sterol, amino acid) biosynthesis (*Lyssiotis and Cantley, 2014*). When this metabolite is in abundance, perhaps it is sensed by the machinery that stimulates TOR signaling, a central regulator of cellular metabolism (*Perera and Zoncu, 2016*). With this in mind, it is worth noting that wortmannin has also been proposed to inhibit PI kinase-related TOR kinases (*Cameroni et al., 2006*). However, activation – not inhibition – of TOR kinase by ER stress stimulates vacuole fission (*Stauffer and Powers, 2015*), and inhibitors of TOR signaling, such as rapamycin, block vacuole fission (*Machiallat et al., 2012*). Thus, it is unlikely that wortmannin or acetate inhibits TOR kinase to stimulate vacuole fission in our preparations. Rather, we suspect that acetate instead blocks PI4-kinase activity preventing genesis of PI-4-P (and subsequently PI-4,5-P_2_) to prevent recruitment and/or stabilization of fusion proteins on vacuole membranes (*Stroupe et al., 2006; Mima and Wickner, 2009*; see Figure 4). This interpretation explains how fusion may be inhibited but how is vacuole fission stimulated? We speculate that by blocking PI-4-P genesis acetate, a larger pool of PI becomes available to be converted into PI-3,5-P_2_ by Vps34 and Fab1. Shunting PI into this pathway would permit the large increase in [PI-3,5-P_2_] necessary to support fission (*Bonagelino et al., 2002*).

**Figure 4.**
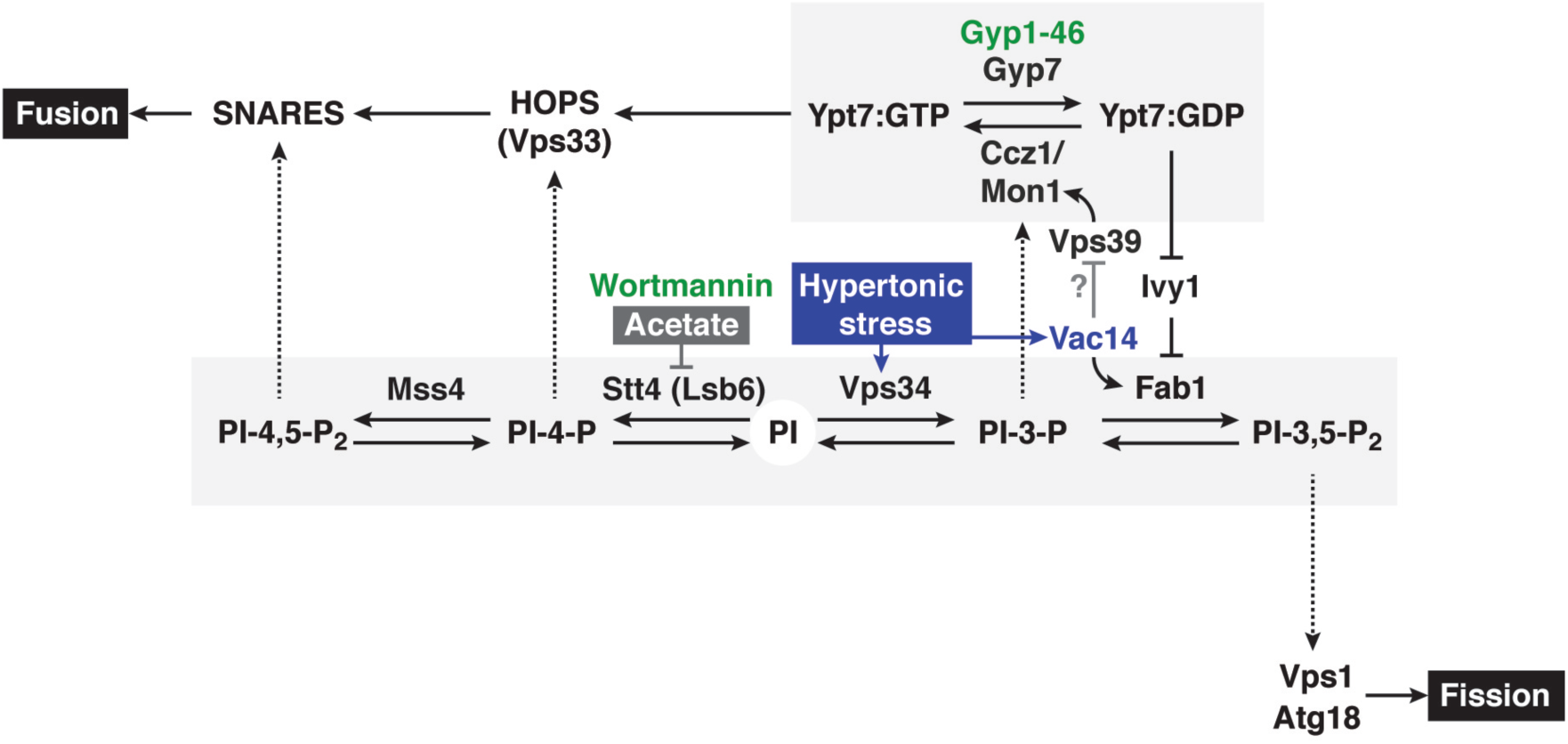
Revised working model of vacuole morphology affected by acetate or hypertonic stress. Diagram depicting molecular interactions between Rab-GTPase and phosphatidylinositol signaling underlying vacuole fission and fusion triggered by acetate or hypertonic stress. Acetate likely inhibits the a PI4-kinase (Stt4 or Lsb6) whereas hypertonic stress primarily targets Rab-GTPase inactivation, possibly through the inhibition of Vps39 by Vac14, to promote vacuole fission over fusion.

### Rab inactivation alone can drive vacuole fission

Previously, hypertonic stress induced by sorbitol was shown to block homotypic vacuole fusion in vitro by inactivating the Rab-GTPase Ypt7 (*Brett and Merz, 2008*). Recently, this mechanism was also shown to promote fission by disrupting the interaction between the I-BAR protein Ivy1 and Ypt7 to activate Fab1 and synthesize PI-3,5-P_2_ (*Malia et al., 2018*). But it remains unclear if lumenal volume loss and subsequent changes in lipid bilayer packing or lateral tension induced by hypertonic stress is necessary for Ivy to activate Fab1.

To assess this possibility, we added rGyp1-46 – a recombinant, purified fragment of the Rab-GTPase Activating Protein (Rab-GAP) Gyp1 that inactivates Ypt7 by promoting GTP hydrolysis (see *Brett and Merz, 2008*) – to vacuole fission reactions containing either KAc, KCl or KCl with 0.8 M glucose (Figure 3B). As expected, increasing concentrations of rGyp1-46 had no effect on fission triggered by hypertonic stress, as this stimulus inactivates Ypt7 to promote fission. Although it had no effect under control, isotonic conditions (125 mM KCl, 200 mM sorbitol), rGyp1-46 stimulated fission in presence of KAc. This important result suggests that (1) unlike hypertonic stress, acetate does not inactivate Ypt7 to promote fission, and (2) inactivating Ypt7 alone is capable of triggering fission. This latter interpretation is consistent with the observations that overexpression of Gyp7, the cognate Rab-GAP for Ypt7, or expression of constitutively inactive YPT7 mutants is sufficient to cause vacuole fragmentation in vivo in the absence of hypertonic stress (*Brett et al., 2008*). Thus, we conclude that Ypt7-inactivation alone is likely sufficient for Ivy1 to activate Fab1.

If not Ivy1, then what senses hypertonic stress? It was previously shown that Vac14, a component of protein complex that includes the PI-3-P5 kinase Fab1, is necessary for Fab1 to be activated by hypertonic stress (*Bonangelino et al., 2002*). From proteomic studies, Vac14 was shown to bind Vps39 (*Elbaz-Alon et al., 2014*), a component of the multisubunit tethering complex HOPS (HOmotypic fusion and Protein Sorting) necessary for Ypt7 activation by Ccz1/Mon1, its cognate Rab-GEF (Guanine nucleotide Exchange Factor; *Nordmann et al., 2010*). If true, this interaction adds an important connection to the existing network underlying this process linking the Fab1-complex to Rab activity (Figure 4). Specifically, we speculate that Vac14 binds and inhibits Vps39 to prevent Ypt7 activation and promote inactivation, presumably by its cognate Rab-GAP Gyp7 (*Brett et al., 2008*). Subsequent release of Ivy1 would stimulate Fab1 (*Malia et al., 2018*). This working model also explains how inactivation of Ypt7 alone triggers fission: By bypassing Vac14 and Vps39, Rab-GAP-mediated inactivation of Ypt7 would simply release Ivy1 to stimulate Fab1 and drive fission. But how then does the vacuole membrane collapse on itself to accommodate membrane scission in the absence of hypertonic stress?

Based on this working model, changes in volume would presumably occur downstream of PI-3,5-P_2_ synthesis (Figure 4). Ion channels, pumps and transporter activities are known to be dependent of their surrounding lipid environments and PI species (e.g. *Hille et al., 2015*). For example, the V-type H^+^-ATPase responsible for maintaining the H^+^-electrochemical gradient is implicated in both vacuole fission and fusion (*Baars et al., 2007; Takeda et al., 2008; Strasser et al., 2011*). Its assembly and stability is presumably enhanced by PI-3,5-P_2_ (*Li et al., 2014*) but not necessarily its activity (*Ho et al., 2015*). This H^+^gradient provides energy to secondary transporters that translocate osmolytes across the vacuole membrane (*Li and Kane, 2009*). Greg Odorizzi’s group recently discovered that Vnx1, a vacuolar Na^+^(K^+^)/H^+^exchanger responsible for lumenal monovalent cation import, is stimulated by deletion of FAB1 (*Wilson et al., 2018*). Moreover, they find that loss of FAB1 blocks TrpY1/Yvc1, a cation channel responsible for lumenal Na^+^, K^+^and Ca^2+^efflux. This result is consistent with TrpY1 being activated by PI-3-P conversion to PI-3,5-P_2_ (*Hamamoto et al., 2018*). Thus, we speculate that PI-3,5-P_2_ promotes net lumenal cation efflux by stimulating TrpY1 and inhibiting Vnx1. Together with vacuolar anion transporters/channels (e.g. the Na^+^/inorganic phosphate symporter Pho89) and aquaporins (that remain unknown), this would create an outward osmotic gradient across the vacuole membrane and drive water efflux to decrease lumenal volume necessary for membrane scission (*Li and Kane, 2009*). Furthermore, this could potentially drive a positive feedback loop through activation of the proposed osmo-sensor Vac14 and Vps39 inhibition, which in turn would further inactivate Ypt7, displace more Ivy1 and drive more PI-3,5-P_2_ synthesis by Fab1.

## Conclusions

In sum, using a new in vitro assay to measure vacuole membrane fission, we refined the existing model of the molecular circuity underlying vacuole morphology, by discovering that acetate triggers fission by blocking PI4-kinase activity whereas hypertonic stress triggers fission by stimulating PI-3-P5-kinase activity through Rab-GTPase inactivation (Figure 4). This infers that PI-4-P and PI-3-P signaling is highly integrated and we speculate that perhaps blocking genesis of PI-4-P with acetate or wortmannin, increases the free pool of PI necessary for synthesis of PI-3-P and subsequently PI-3,5-P_2_ needed for fission (*Weisman, 2003*). We also confirm that these stimuli have the opposing effect on homotypic vacuole fusion, lending additional support to the idea that fission and fusion are highly coordinated (e.g. *Alpadi et al., 2013*). This is a requirement for efficient changes in morphology, whereby if fission occurs, the counteracting process of fusion must be blocked for the organelle to remain fragmented, i.e. decrease size and increase number (*LaGrassa and Ungermann, 2005*). Thus, it is not surprising that PI and Rab-GTPase signaling, known to be critical for fusion, also mediate fission. Because vacuoles do not need to be stained with a fluorescent dye and small reaction volumes are microplate compatible, this new fission assay can be easily scaled up to accommodate high-content screening experiments. Thus, it sets the stage for future studies that will test this revised model and further reveal the complex molecular interactions underlying organelle fission and morphology in molecular detail.

## MATERIALS AND METHODS

### Yeast strains and reagents

We used the *S. cerevisiae* strain SEY6210 *pep4*Δ Vph1-GFP [*MATα leu2-3 ura3-52 his3-Δ200 trp1-Δ901 suc2-Δ9 lys2-801 pep4∷HIS3 VPH1-GFP (TRP1)*] for the fluorescence-based in vitro fission assay and vacuole membrane detection by fluorescence microscopy (see *McNally et al., 2017*). BY4742 *pep4*Δ (*MATα leu2-3 ura3-52 his3-Δ200 lys2-801 pep4∷NEO)* or *pho8*Δ (*MATα leu2-3 ura3-52 his3-Δ200 lys2-801 pho8∷NEO) S. cerevisiae* strains purchased from Invitrogen (Carlsbad, CA, USA) were used for the in vitro fusion assay. All yeast growth media was purchased from BioShop Inc. (Burlington, ON, Canada). Buffer ingredients and reagents were purchased from Sigma Aldrich (St. Louis, MI, USA) with the exception of ficoll from GE Healthcare (Tokyo, Japan) and ATP from Roche (Indianapolis, IN, US). Recombinant Gyp1-46 protein and oxalyticase were expressed in *E.coli* and purified by affinity chromatography as previously described (*see Karim et al., 2018*). All proteins or reagents added to in vitro fusion or fission reactions were diluted in or buffer exchanged into PS buffer (20 mM PIPES, 200 mM sorbitol), aliquoted, flash frozen in liquid nitrogen and stored at –80 °C until use.

### Yeast vacuole isolation

Yeast cultures were grown in a shaking incubator overnight at 30 °C in 1 L YPD medium to a density of 1.4 – 1.8 OD600_nm_/mL. Cells were then harvested by centrifugation (3,000 g for 10 minutes at 4 °C), washed (10 minutes at 30 °C) with 50 mL buffer containing 100 µM DTT and 50 mM Tris-HCl pH 9.4, sedimented (3, 500 g for 5 minutes at room temperature), resuspended in 15 mL spheroplasting buffer (25 mM potassium phosphate pH 6.8 and 200 mM sorbitol in 1:20 YPD medium diluted in water) containing 1 – 2 µg/mL purified oxalyticase, and incubated for 30 minutes at 30 °C. Spheroplasts were collected by centrifugation (1,250 g for 2 minutes at 4 °C), resuspended in 2 mL ice-cold PS buffer (20 mM PIPES, 200 mM sorbitol) containing 15 % ficoll, and treated with 0.2 – 0.4 µg/mL DEAE dextran for 3 minutes at 30 °C to disrupt the plasma membrane. Permeabilized spheroplasts were then transferred to an ultracentrifuge tube on ice, 8 %, 4 % and 0 % ficoll layers were added on top, and samples were subjected to high-speed centrifugation (125,000 g for 90 minutes at 4 °C) to isolate vacuoles from other cell components. Vacuoles were then collected from interface between 4 and 0 % ficoll layers and placed on ice until use. Vacuole protein concentrations were determined by Bradford assay.

### In vitro vacuole fission assay

To quantify vacuole membrane fission in vitro, we prepared 30 µL fission reactions by adding 6 µg of vacuoles isolated from SEY6210 *pep4*Δ Vph1-GFP cells to standard fission reaction buffer (PS buffer containing 5 mM MgCl_2_, 125 mM KCl, 10 mM CoA, and 1 mM ATP to stimulate fission; *see Michaillat et al., 2012*) and then incubated them at 27 °C for 30 minutes. Where indicated, increasing concentrations of glucose, sorbitol, wortmannin or recombinant Gyp1-46 protein were added, or KAc was replaced with KCl, prior to incubation. Reactions were then subjected to centrifugation (3,000 g for 3 minutes at 4 °C) to separate small vacuoles (present in the supernatant) from larger vacuoles (present in the pellet). Of note, to possibly improve isolation of fission products, we added increasing amounts of trypsin after fission reactions were completed to cleave tethering proteins that may keep newly formed fragments attached to precursor organelles. But trypsin incubation had no discernable effect on fission product isolation by centrifugation (data not shown). After collecting the supernatant, pellets were resuspended in 20 µL fission reaction buffer and both samples were then transferred to a black conical-bottom 96-well microplate. GFP fluorescence (λ_ex_ = 485 nm, λ_em_ = 520 nm) was then measured using a Synergy H1 multimode microplate reader (BioTek Instruments Inc., Winooski, VT, USA), values were background subtracted and the ration of supernatant over pellet fluorescence was calculated as a measure of vacuole membrane fission in vitro. Data shown was normalized to the value obtained under control (no treatment), isotonic conditions. Reaction buffer osmolarity was confirmed using a Vapro 5520 vapor-pressure osmometer (Wescor, Logan, UT, USA). Vacuole diameter was measured using a Brookhaven 90 Plus Particle Size Analyzer (Brookhaven Instruments Cooperation).

### In vitro homotypic vacuole fusion assay

Homotypic vacuole fusion in vitro was measured using a colorimetric assay that relies on maturation of the alkaline phosphatase Pho8 (*see Brett and Merz, 2008*). In brief, 30 µL fusion reactions were prepared by adding 3 µg of vacuoles isolated from BY4742 *pho8*Δ cells and 3 µg of vacuoles isolated form BY4742 *pep4*Δ cells to standard fusion reaction buffer (PS buffer containing 125 mM KCl, 5 mM MgCl_2_, 10 µM CoA and 1 mM ATP to simulate fusion) and then incubated at 27 °C for 90 minutes. Where indicated increasing concentrations of sorbitol were added or KAc was replaced with KCl, prior to incubation. Upon membrane fusion, lumenal content mixing permits immature Pho8 (within vacuoles from cells missing Pep4) to be cleaved by the protease Pep4 (within vacuoles from cells missing Pho8) to activate the enzyme. Pho8 activity is then measured by adding 500 µL development buffer (250 mM Tris-HCl pH 8.5, 10 mM MgCl_2_, 0.4 % triton X-100) containing 1 mM paranitrophenolphosphate, a Pho8 substrate, and incubated for 5 minutes at 30 °C. The phosphatase reaction was terminated with 500 µL stop buffer (100 mM glycine pH 11) and the absorbance of the yellow product, paranitrophenol, was measured at 400 nm using a NanoDrop 2000c spectrophotometer (Thermo Fisher Scientific, Waltham, MA, USA). A_400nm_ values were background subtracted and normalized to values obtained under control, isotonic conditions (125 mM KCl).

### Fluorescence microscopy

Using HILO (Highly Inclined and Laminated Optical sheet) microscopy, images of fission reactions containing vacuoles isolated from SEY6210 *pep4*Δ Vph1-GFP cells were acquired using a Nikon Eclipse TiE inverted microscope outfitted with a TIRF (Total Internal Reflection Fluorescence) illumination unit, Photometrics Evolve 512 EM-CCD camera, CFI ApoTIRF 1.49 NA 100x objective lens, and 50 mW 488 nm solid-state laser operated with Nikon Elements software (Nikon Canada Inc., Mississauga, ON, Canada). Images were acquired 1 µm into the sample. Micrographs shown were adjusted for brightness and contrast, inverted and sharpened with an unsharpen masking filter using Image J (National Institutes of Health, Bethesda, MD, USA) and Photoshop CC software (Adobe Systems, San Jose, CA, USA).

### Data analysis and presentation

All quantitative data was processed using Microsoft Excel software (Microsoft Corp., Redmond, WA, USA). Data was plotted using Kaleida Graph v.4.0 software (Synergy Software, Reading, PA, USA) and figure panels were prepared using Illustrator CC software (Adobe Systems, San Jose, CA, USA). Means ± S.E.M. are shown and Student’s two-tailed t-tests were used to assess significance (*P < 0.05). Micrographs shown are best representatives of 5 biological replicates (each replicate represents a sample prepared from a separate yeast culture on different days), imaged at least 5 times each (technical replicates) whereby each field examined contained > 83 vacuoles. Fission and fusion data shown represent 3 or more biological replicates (each replicate represents a sample prepared from a separate yeast culture on different days) conducted in duplicate (technical replicates).

## AUTHOR CONTRIBUTIONS

D.P. and C.L.B conceived the project, performed experiments, and prepared data for publication. C.L.B. wrote the paper.

## ACKNOWLEDGEMENTS

We thank Andrew Chapman for use of his osmometer and Jack Kornblatt for use of his particle size analyzer. This work was supported by Natural Sciences and Engineering Research Council of Canada grants RGPIN/403537-2011 and RGPIN/2017-06652 to C.L.B.

**ABBREVIATIONS**
CMOS: complementary metal-oxide semiconductor
EMCCD: Electron Multiplying Charge Coupled Device
ER: endoplasmic reticulum
GAP: GTPase Activating Protein
GEF: Guanine nucleotide Exchange Factor
GFP: green fluorescent protein
HOPS: homotypic fusion and protein sorting
I-BAR: inverted BAR (Bin, Amphiphysin, Rvs)
PI: phosphatidylinositol
PI-3-P: phosphatidylinositol-3-phosphate
PI-3,5-P: phosphatidylinositol-3,5-diphosphate
PI-4-P: phosphatidylinositol-4-phosphate
PI-4,5-P: phosphatidylinositol-4,5-diphosphate
PS buffer: 20 mM PIPES, 200 mM sorbitol buffer
ROI: region of interest
SEM: standard error of the mean
TOR: target of rapamycin
VPS: vacuole protein sorting

